# Hydrogel formulation controls size-dependent accumulation of proteins and nanoparticles in PEG microparticles

**DOI:** 10.64898/2026.08.07.741917

**Authors:** Alyssa Arnheim, Ian Morales, Andrew Tran, Dino Di Carlo

## Abstract

Hydrogels are widely used in sensing, delivery, and tissue engineering because their transport properties can be tuned through material design. However, while hydrogel permeability is often characterized using small molecules, many practical applications depend on the uptake and retention of much larger species, including protein conjugates and nanoparticles. Here, we systematically investigate how polyethylene glycol (PEG)-acrylate hydrogel microparticle formulation influences accumulation of signal-generating probes spanning a broad size range. We fabricated particles across a 36-condition design space varying nominal PEG-acrylate molecular weight, polymer weight percent, and UV crosslinking dose, and related formulation-dependent probe accumulation to particle swelling behavior. Increasing nominal PEG-acrylate molecular weight and decreasing polymer weight percent produced more highly swollen particles and strongly enhanced accumulation of fluorescent streptavidin conjugates, with the largest effects observed for bulky labels such as allophycocyanin and phycoerythrin. Gold nanoparticle accumulation was even more formulation-restricted, with detectable colorimetric signal observed primarily in the most permissive formulations. These findings establish design rules linking PEG hydrogel formulation to size-dependent accumulation and show that formulations suitable for small probes may be inadequate for larger reporters. More broadly, this framework may inform the design of hydrogels for particle-based assays as well as other applications where transport of macromolecules or nanoscale materials is important.

## Introduction

Hydrogels are widely used in biomedical engineering because they combine biocompatibility with tunable mechanical and transport properties.^1–3^ Their highly hydrated polymer networks can be engineered to control stiffness,^4^ swelling,^5^ permeability,^6–8^ and molecular retention,^9^ making them useful in applications ranging from contact lenses^10,11^ and wound dressings^12,13^ to tissue scaffolds,^14–16^ drug delivery depots,^10,17^ biosensors,^18,19^ and diagnostic materials.^20–23^ In research settings, hydrogels are also increasingly used as defined microenvironments for cell culture,^24,25^ organoid growth,^26–28^ and assay compartmentalization.^29,30^ Across these diverse uses, the ability of a hydrogel to admit, exclude, retain, or release molecules is often central to its performance.

This transport behavior is particularly important when hydrogels are used as functional assay materials rather than passive structural supports. In particle-based sensing and lab-on-a-particle systems, for example, assay performance depends on whether analytes, affinity reagents, and reporting moieties can access the interior of the hydrogel and accumulate there at sufficient levels for detection.^20,21,31–35^ This requirement is not limited to fluorescence-based assays. Reliable signal generation may require local enrichment of fluorophore-labeled proteins, reporters, enzymes, chromogenic substrates, or nanoparticle labels such as gold nanoparticles.^36–38^ If these components cannot efficiently penetrate or be retained within the network, signal remains surface bound and weak even when binding chemistry is otherwise favorable.

Although hydrogel transport has been studied extensively, much of that work has focused on small-molecule diffusion or bulk release behavior.^3,17,39,40^ Though studies exist, comparatively less is known about how hydrogel formulation controls the uptake and accumulation of larger species spanning protein conjugates and nanoparticles.^41–45^ This is an important gap because many practical readout strategies rely on labels that are substantially larger than the small molecules often used to characterize hydrogel permeability. As a result, design principles optimized for one class of cargo may not translate across all sizes.

Here, we use PEG-acrylate hydrogel microparticles as a model synthetic system to systematically investigate how network-forming parameters influence size-dependent accumulation. PEG-based hydrogels are well suited for this purpose because their structure can be tuned reproducibly through polymer molecular weight, polymer weight fraction, and crosslinking conditions. We therefore varied nominal PEG-acrylate molecular weight, PEG weight percent, and UV crosslinking conditions to generate a panel of particle formulations and measured their ability to accumulate signal from probes spanning a wide size range, including fluorescent protein conjugates and gold nanoparticles. We further relate these trends to particle swelling behavior as a practical readout of network expansion.

Our goal is to establish formulation-level design rules for hydrogel particles that support the accumulation of large signal-generating species. More broadly, this framework connects hydrogel structure to functional uptake behavior in a way that may inform not only particle-based sensing systems, but also other hydrogel applications in drug delivery and tissue engineering where transport of larger biomolecules or nanoscale materials into or out of hydrogels is important.

## Results

### Particle fabrication and swelling behavior

We fabricated PEG-acrylate hydrogel microparticles using droplet microfluidics and UV crosslinking, enabling systematic variation of hydrogel formulation across a 36-condition design space (Figure 1a-c). Formulations were generated by varying nominal PEG-acrylate molecular weight (5–40 kDa), PEG weight percent (7.5–22.5 wt%), and UV dose (37–172 mJ/cm^2^). We selected these parameters because each is expected to influence hydrogel network formation: increasing nominal PEG-acrylate molecular weight and decreasing polymer weight percent should reduce crosslink density and promote a more expanded network, whereas UV dose may modulate the extent of polymerization. This formulation panel provided a controlled framework for testing how hydrogel design affects the accumulation of probes spanning a wide size range.

**Figure 1.**
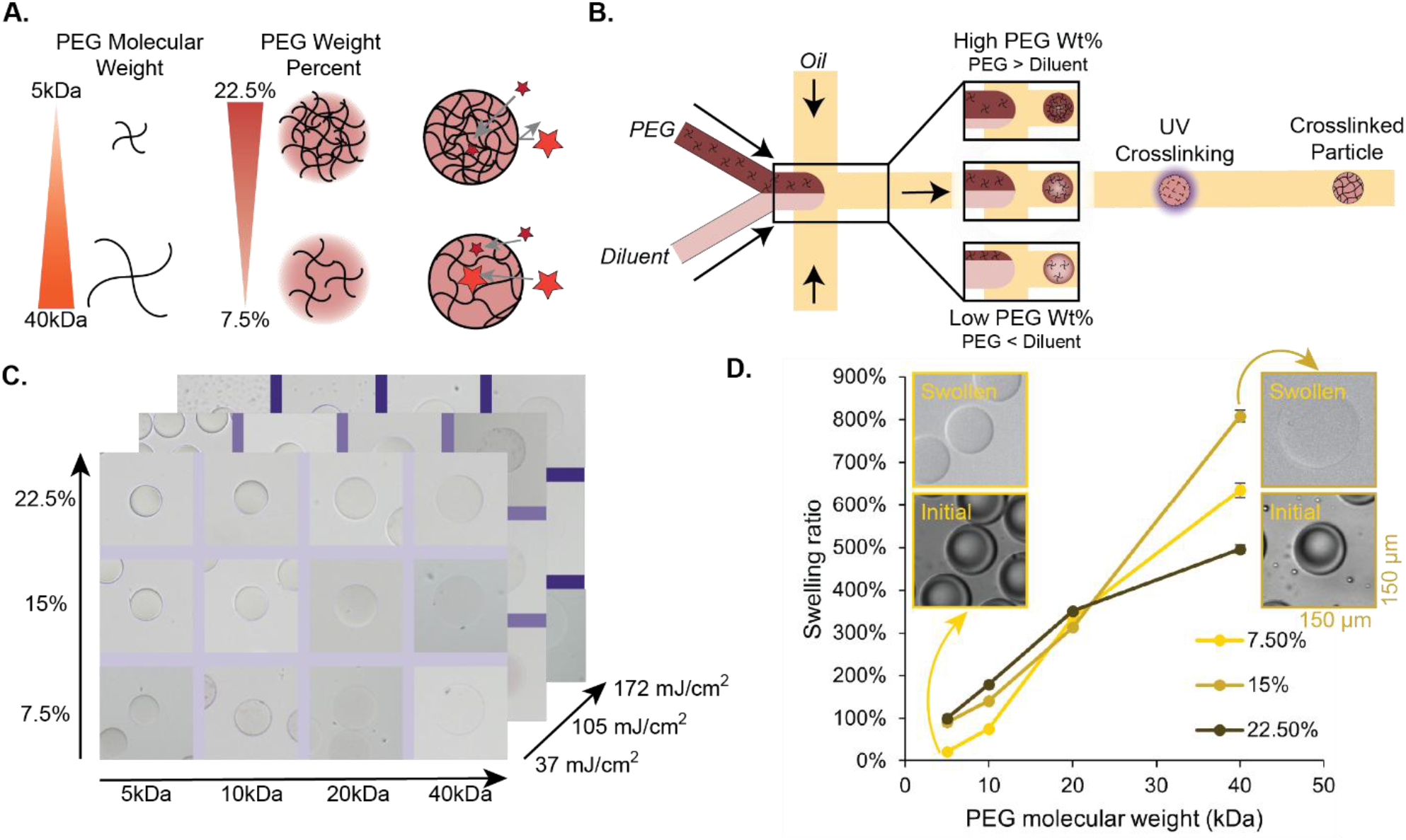
Tuning PEG hydrogel formulation to enable size-dependent signal accumulation. (A) Schematic of how changes in formulation (changing 4-arm PEG acrylate MW and PEG wt%) are expected to impact final particle mesh size. (B) Droplet microfluidic fabrication workflow and UV crosslinking. (C) Parameter space explored (PEG arm MW, PEG wt%, and UV dose). (D) Swelling ratio vs PEG molecular weight for different weight % formulations.

To assess how formulation altered bulk hydrogel expansion, we measured particle swelling ratio across formulations (Figure 1d). Swelling increased strongly with nominal PEG-acrylate molecular weight and decreased with PEG weight percent, which is consistent with these parameters producing more expanded hydrogel networks. The largest effect was observed at 7.5 wt%, where swelling increased from 22% for 5 kDa PEG to 635% for 40 kDa PEG. This trend was preserved within each weight-percent series, indicating that nominal PEG-acrylate molecular weight was a dominant determinant of swelling behavior. By contrast, changes in UV dose produced comparatively small and inconsistent effects across formulations (Supplementary Fig. 1). Together, these results establish nominal PEG-acrylate molecular weight and PEG weight percent as the principal formulation variables controlling particle expansion.

### Expanded hydrogel formulations support greater accumulation of fluorescent protein probes

To test whether the formulation-dependent differences in particle swelling translated into functional differences in probe uptake, we incubated biotinylated hydrogel particles with streptavidin-conjugated fluorescent probes and measured particle-associated fluorescence after washing (Figure 2a). This assay was designed to evaluate whether hydrogel formulations that form more expanded networks are better able to admit and retain bulky fluorescent labels that are commonly used in biomolecular assays. We examined three streptavidin conjugates spanning a range of reporter sizes: streptavidin–Alexa Fluor 555 (∼60 kDa), streptavidin–allophycocyanin (APC, ∼165 kDa), and streptavidin–phycoerythrin (PE, ∼300 kDa) (Figure 2c).

**Figure 2.**
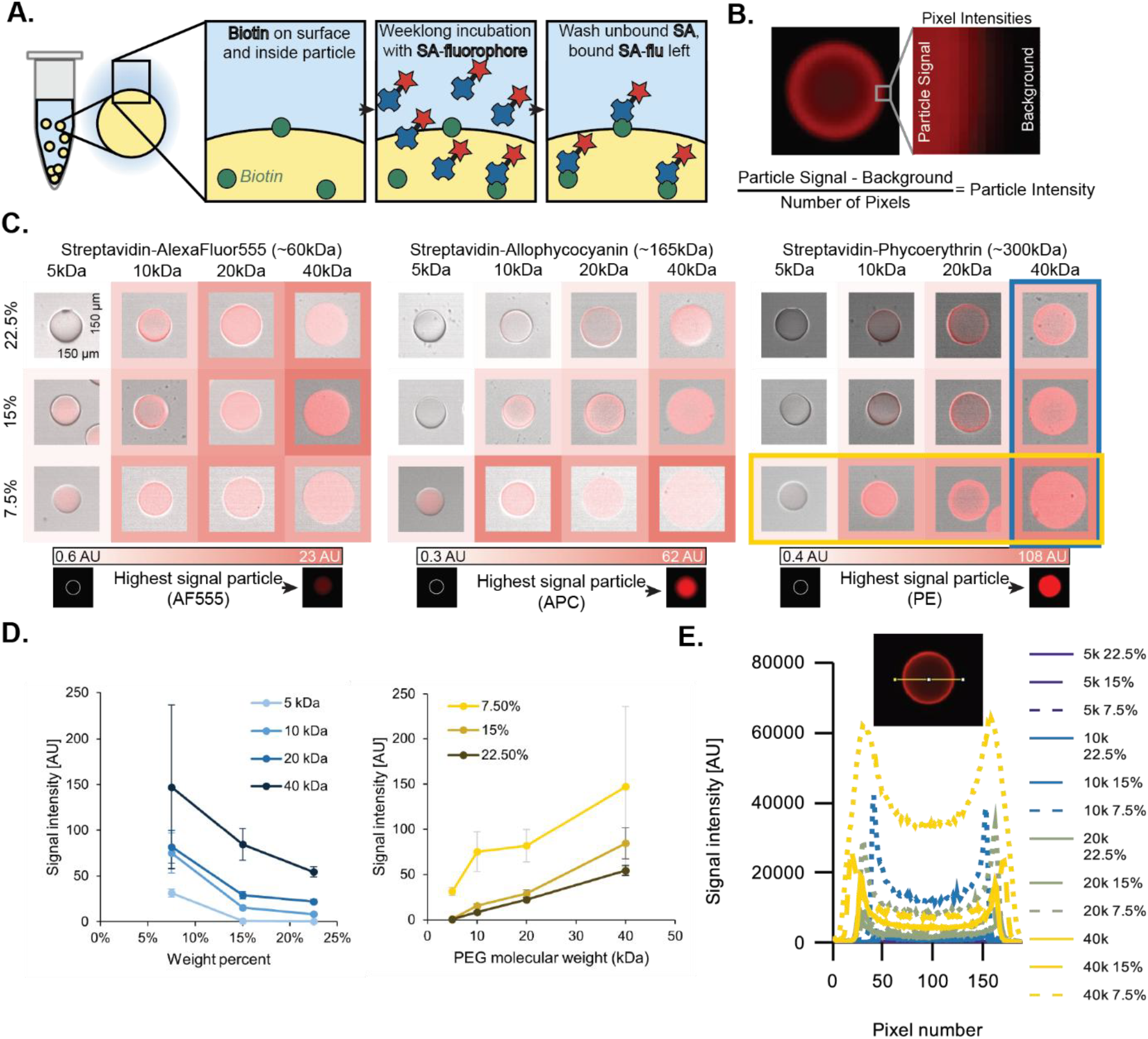
Fluorescent probe accumulation increases with nominal PEG-acrylate molecular weight and decreases with PEG weight percent. (A) Schematic of the fluorescent probe accumulation assay. Biotinylated hydrogel particles were incubated with streptavidin-conjugated fluorophores, washed, and analyzed for particle-associated fluorescence. A one-week incubation was selected to provide an extended and standardized period for probe access and accumulation across formulations. (Supplementary Fig. 4) (B) Fluorescent image of SA-PE penetration into a hydrogel particle. Inset demonstrates analysis metric used for particle-associated signal. (C) Representative fluorescence images and quantified mean background-subtracted fluorescence normalized by particle area for streptavidin–Alexa Fluor 555, streptavidin–APC, and streptavidin–PE across nominal PEG-acrylate molecular weight and PEG weight percent formulations. Each particle is pictured within a 150µm x 150 µm box. Heatmaps summarize within-probe formulation dependence. Color scales are normalized within each probe, with the fluorescence images below the gradients demonstrating highest signal particles for each probe. (D) Quantification of PE accumulation showing increased particle-associated fluorescence with increasing nominal PEG-acrylate molecular weight and decreasing PEG weight percent. (E) Fluorescence cross-section intensity profiles for representative particle formulations plotted on a common intensity scale, illustrating stronger and more spatially extensive fluorescence in high-molecular-weight, low-weight-percent formulations.

Across all three probes, representative fluorescence images revealed a consistent formulation dependence: particle-associated signal increased with increasing nominal PEG-acrylate molecular weight and decreasing PEG weight percent (Figure 2c). This trend became progressively more pronounced for the larger fluorescent conjugates. Particles formed from low-molecular-weight PEG and high polymer weight percent showed weak fluorescence, whereas particles fabricated from high-molecular-weight PEG at low weight percent showed substantially stronger signal. We quantified these effects by measuring background-subtracted fluorescence normalized by particle area to account for formulation-dependent differences in particle size (Figure 2b, Supplementary Fig. 2).^46^ For PE, which showed the clearest dependence on hydrogel formulation, signal increased monotonically with nominal PEG-acrylate molecular weight within each weight-percent series and decreased as PEG weight percent increased within each molecular-weight series (Figure 2c,d), which is consistent with the swelling measurements in Figure 1d and supporting the interpretation that more expanded hydrogel networks more readily accommodate large protein probes. In contrast, UV dose did not produce a consistent trend in fluorescence intensity across formulations or probes: increasing UV dose sometimes increased and sometimes decreased particle-associated signal, and no monotonic UV-dependent behavior was apparent relative to the stronger effects of nominal PEG-acrylate molecular weight and PEG weight percent (Supplementary Fig. 3).

To compare how fluorescence was spatially distributed within particles across formulations, we plotted particle intensity profiles on a common scale (Figure 2e). In addition to the overall increase in signal for the more permissive formulations, the profiles revealed a recurring edge-enhanced pattern, with fluorescence maxima near the particle periphery rather than at the center. For a spherical particle imaged through its full depth, one would typically expect the greatest integrated intensity at the center because the optical path length is largest there. The observed peripheral peaks therefore suggest that probe accumulation was enriched near the particle boundary. One possible explanation is that the particle edge is structurally distinct from the core. For example, oxygen inhibition during UV crosslinking reduces crosslinking efficiency near the droplet interface and produces a more permissive outer region. Alternatively, the pattern may reflect transport-related effects, such as preferential loading or retention near the particle surface. Regardless of mechanism, these profiles indicate that differences between formulations reflected broader and spatially structured changes in particle-localized fluorescence.

These results show that hydrogel formulation strongly controls the accumulation of fluorescent protein probes and that this dependence becomes more important as probe size increases. Formulations composed of higher-molecular-weight PEG and lower polymer weight percent consistently supported the greatest accumulation, indicating that hydrogel particles optimized for small labels may not be optimal for larger, brighter fluorescent reporters such as APC and PE.

### Gold nanoparticle accumulation is restricted to the most permissive formulations

To test whether the same formulation trends extended to even larger signal-generating species, we next examined the accumulation of streptavidin-conjugated gold nanoparticles (AuNPs) with diameters of 5, 10, and 20 nm. These particles represent a substantially more demanding transport case than the fluorescent probes in Figure 2 because of their larger size and because detectable colorimetric readout requires sufficient local nanoparticle loading within the hydrogel. We therefore focused on whether hydrogel formulations that supported strong accumulation of large fluorescent probes would also permit entry and visible retention of AuNPs.

Across nanoparticle sizes, visible particle-associated gold signal was strongly formulation-dependent (Figure 3). For all three AuNP diameters, detectable accumulation above background was observed primarily in particles fabricated from high-molecular-weight PEG at low polymer weight percent, with the strongest signal occurring in the 40 kDa, 7.5 wt% formulation. In these particles, AuNP loading appeared as a pink haze distributed within the particle interior. By contrast, formulations made from lower-molecular-weight PEG and/or higher polymer weight percent showed little to no visible signal, consistent with a more restrictive hydrogel network that limited nanoparticle entry or retained loading below the threshold required for optical detection.

**Figure 3.**
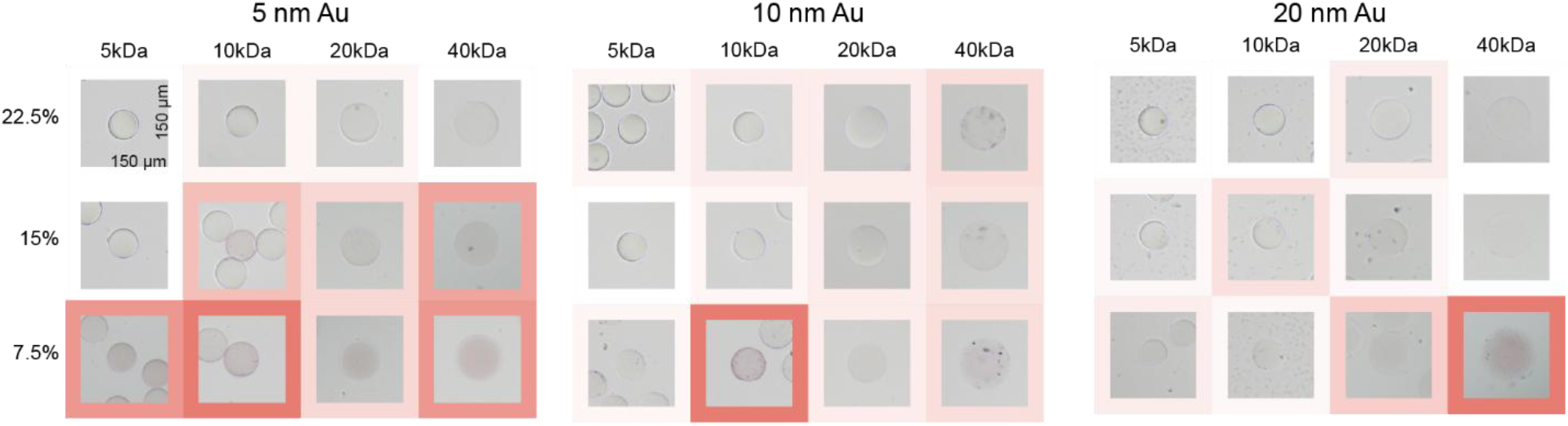
Gold nanoparticle accumulation is restricted to high-molecular-weight, low-weight-percent hydrogel formulations. Representative brightfield images of particles incubated with streptavidin-conjugated 5, 10, or 20 nm gold nanoparticles across nominal PEG-acrylate molecular weight and PEG weight percent formulations. Each particle is pictured within a 150µm x 150 µm box. Particle-associated AuNP accumulation appears as a pink haze within the particle interior. Visibly detectable signal is observed primarily in high-molecular-weight, low-weight-percent formulations, with the strongest accumulation in the 40 kDa, 7.5 wt% condition

The same general formulation dependence was maintained across 5, 10, and 20 nm AuNPs, but the requirement for a permissive network became especially apparent as particle size increased. Whereas some signal could be observed in a small number of intermediate formulations for 5 nm AuNPs, consistent accumulation across nanoparticle sizes was confined to the most expanded formulation space. This result extends the trends observed with APC and PE in Figure 2 and indicates that hydrogel formulation exerts increasingly strong control over accumulation as the reporting species becomes larger and more structurally complex.

Together, these results show that design rules inferred from small or moderately sized probes do not necessarily translate to nanoparticle-based readouts. Instead, colorimetric particle assays that rely on AuNP accumulation require hydrogel formulations with substantially more permissive network structure than those sufficient for smaller fluorescent labels. This finding is particularly relevant for assay design because gold nanoparticles are widely used as robust visual reporters, yet their utility in hydrogel particles depends on whether the material can support enough localized loading to generate a detectable optical signal.

### Nanoparticle-based assay proof of concept

Because the 40 kDa, 7.5 wt% formulation showed the strongest AuNP accumulation across sizes, we selected this high-porosity formulation to test whether enhanced nanoparticle loading could be leveraged in an assay format proof-of-concept. Particles were functionalized with streptavidin, incubated with a titration series of biotinylated goat anti-mouse antibody (0, 1, and 100 µg/mL), and labeled with mouse antibodies conjugated to 5 nm gold nanoparticles (3 OD). Particle-associated color signal increased with increasing biotinylated goat anti-mouse concentration, producing progressively stronger visible coloration and higher measured color saturation (Figure 4).

**Figure 4.**
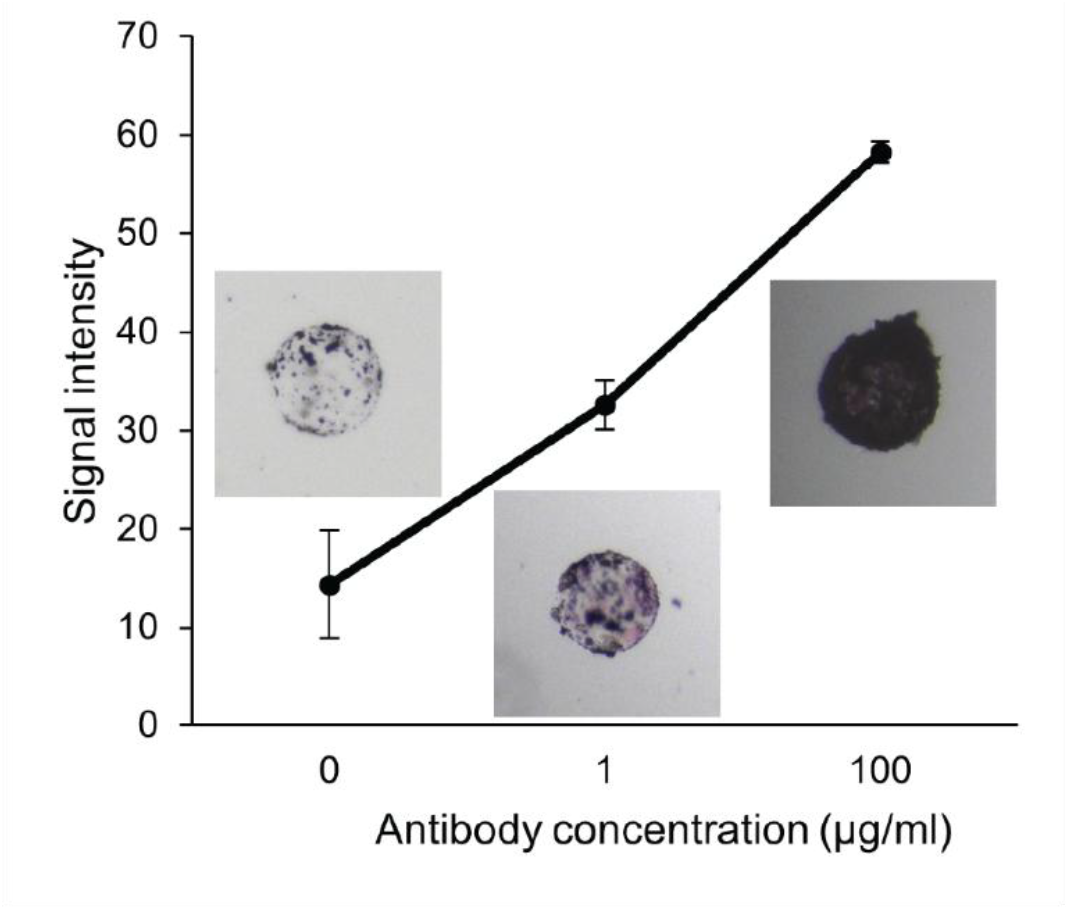
Gold-based immunoassay-style labeling on high-porosity hydrogel particles produces tunable colorimetric signal. Each particle is pictured within a 170µm x 170 µm box. Capture-antibody titration performed on the highest-porosity formulation (40 kDa, 7.5 wt%). Particles were functionalized with streptavidin, incubated with biotinylated goat anti-mouse capture antibody (0– 100 µg/mL), and labeled with mouse antibody–gold (5 nm, 3 OD). Increasing capture antibody concentration increased visible particle coloration and HSV saturation quantified within a particle mask; the 0 µg/mL condition served as the negative control.

Color intensity was quantified by converting images to HSV color space and calculating the mean saturation within a particle mask. The 0 µg/mL biotinylated goat anti-mouse condition served as the negative control; in the absence of capture antibody, any residual particle-associated signal reflects baseline/nonspecific accumulation of the gold-labeled antibody.

Increasing immobilized antibody concentration produced a corresponding increase in particle-localized gold signal, demonstrating that the permissive formulation can support affinity-dependent accumulation of gold-conjugated reagents.

## Discussion

While hydrogel transport has been studied extensively, much of the prior characterization emphasizes small-molecule diffusion and bulk release behavior, leaving comparatively less clarity on how formulation governs uptake and accumulation of larger species (protein conjugates and nanoparticles) in functional assay contexts.^1,2,44,45^ In hydrogel microparticles specifically, this gap is important to investigate because assay performance may depend on both binding chemistry and whether reporters can access the interior of the particle and remain there at detectable levels.

By sweeping a 36-condition PEG-acrylate formulation space, we identify composition-driven design rules that dominate accumulation behavior. Increasing nominal PEG-acrylate molecular weight (5–40 kDa) and decreasing PEG weight percent (7.5–22.5 wt%) produced markedly higher swelling and correspondingly higher probe accumulation, consistent with reduced crosslink density and a more expanded, permissive network. UV dose (37–172 mJ/cm^2^) produced inconsistent effects across the panel, and UV-dependent differences were difficult to resolve relative to the clear effects of PEG molecular weight and wt%. This suggests that, within the dose range tested, network “openness” is controlled primarily through bulk formulation rather than UV dose alone.

A key mechanistic insight from the fluorescence experiments is that formulation affects both how much signal accumulates, and where it accumulates within the particle. The recurrent edge-enhanced intensity profiles indicate preferential enrichment near the periphery rather than uniform volumetric loading. One plausible explanation is an interface-driven structural gradient (e.g. oxygen inhibition reducing crosslinking efficiency near the droplet boundary),^47^ though transport-limited loading or retention near the surface could also contribute. From an assay-design perspective, this matters because “particle-associated signal” may reflect a mix of true interior permeability and surface-adjacent enrichment—both of which can be useful for readout but may behave differently as reporter size increases.

Unlike fluorescent probe accumulation, which has intrinsically higher signal to noise ratio given the dark background, gold nanoparticles as signal generators were shown to require more stringent conditions to enable colorimetric lab-on-a-particle assays. This is because development of signal requires sufficient particle-localized accumulation to cross a detection threshold with weakly scattering particles. With that added constraint, detectable particle-associated signal across 5, 10, and 20 nm AuNPs was largely restricted to the most expanded formulations particularly the 40 kDa, 7.5 wt% condition. This result reinforces that design rules inferred from smaller probes may not translate to nanoparticle-based readouts, and it provides a concrete formulation target for enabling colorimetric reporters in hydrogel particles.

Importantly, the gold accumulation results translate naturally into a point-of-care-relevant immunoassay stack. Because the 40 kDa, 7.5 wt% formulation showed the strongest AuNP accumulation, it was selected for an assay-format demonstration, and particle color increased with increasing capture antibody concentration when binding to AuNP-labeled targets. Quantification via saturation within a particle mask provides a straightforward path to instrumented or semi-instrumented readout (e.g., microscopy/cytometry), while also utilizing well-developed infrastructure of gold-nanoparticle colorimetric reagents used in many rapid tests. These results suggest a route toward particle-based colorimetric diagnostics where numerous hydrogel particles can act as the accumulation scaffolds for gold labeling.

Beyond assay performance, this work expands general understanding of PEG hydrogel behavior at the extremes explored here. We identified conditions that enable crosslinking that maintains structural integrity in highly permissive networks (e.g., 40 kDa PEG at low wt%). In our hands, formulations below ∼5 wt% did not crosslink reliably (not shown), defining a practical lower boundary for pursuing even larger mesh sizes in this chemistry. These “fabrication limits” provide a helpful design space for materials scientists and engineers looking to use highly permeable microgels.

Finally, the implications extend beyond diagnostics. Highly permissive particle formulations that support transport of large protein probes suggest a materials toolkit for macroporous scaffolds assembled from microparticles (e.g., microporous annealed particle-style materials):^48–50^ if individual building-block particles do not substantially hinder macromolecular transport, assembled scaffolds can be designed with predictable accessibility for proteins and other large cues. The observed formulation-dependent permeability also raises the possibility of heterogeneous MAP scaffolds, where mixing particles with different polymer network properties or functional densities yields composite materials with spatially tuned transport, retention, or signaling behavior. Overall, by selecting formulations according to the size and chemistry of the species that must enter, accumulate, or be retained, the design rules reported here offer a foundation for creating tunable hydrogel microparticle systems that span from particle-based colorimetric immunoassays to engineered macroporous culture environments.

## Methods

### Particle fabrication

Biotinylated PEG hydrogel microparticles (∼50 µm radius) were fabricated using a three-inlet flow-focusing microfluidic device cast in polydimethylsiloxane (PDMS). Devices were made by curing PDMS on an SU-8 master mold and plasma bonding to a glass slide.^51^ Two aqueous streams and one oil stream were used to generate monodisperse water-in-oil droplets that were subsequently crosslinked by UV illumination. Flow was controlled using syringe pumps (Harvard Apparatus PHD 2000).

Aqueous inlet 1 (PEG prepolymer stock). The PEG prepolymer stream contained 25% (w/v) four-arm PEG-acrylate (Advanced BioChemicals) dissolved in PBS, with PEG-acrylate molecular weight varied across experiments (5, 10, 20, or 40 kDa). Lithium phenyl-2,4,6-trimethylbenzoylphosphinate (LAP; Sigma) was included at 2.5% (w/v), and acrylate-PEG-biotin (5 kDa; Nanocs) was included at 10% (v/v).

Aqueous inlet 2 (diluent/initiator stock). The diluent stream contained 2.5% (w/v) LAP and 10% (v/v) acrylate-PEG-biotin (5 kDa) in PBS and did not contain 4-arm PEG-acrylate. This stream enabled dilution of PEG-acrylate while maintaining constant LAP and acrylate-PEG-biotin concentrations across formulations. Because both aqueous inlets contained LAP and acrylate-PEG-biotin at the same concentrations, the final concentrations in droplets were 2.5% (w/v) LAP and 10% (v/v) acrylate-PEG-biotin for all conditions.

Oil phase. The continuous oil phase consisted of 0.5% (v/v) Pico-Surf (Sphere Fluidics) in Novec 7500.

Droplets were polymerized under focused UV illumination through a DAPI filter set and microscope objective (Nikon Eclipse Ti-S) using a 10× objective. UV crosslinking was performed using a constant exposure time of 0.5 s at three dose conditions: 37, 105, and 172 mJ/cm^2^.

### Tuning PEG wt% by on-chip dilution

PEG wt% in the final droplets was tuned by changing the flow-rate ratio of the PEG prepolymer stock (25% w/v PEG-acrylate) and the diluent/initiator stock (0% PEG-acrylate), while keeping total aqueous flow constant (2.5 µL/min) and oil flow constant (15 µL/min). Final PEG-acrylate wt% values were:

- 22.5% PEG: PEG stock 2.25 µL/min + diluent 0.25 µL/min
- 15% PEG: PEG stock 1.5 µL/min + diluent 1.0 µL/min
- 7.5% PEG: PEG stock 0.75 µL/min + diluent 1.75 µL/min

This dilution scheme yields the target PEG wt% because: final PEG wt% = (25% PEG stock) × (PEG stock flow / total aqueous flow).

### Experimental design (36 formulations)

Particles were fabricated across 36 condition combinations comprising:

- PEG-acrylate molecular weight: 5, 10, 20, 40 kDa
- PEG-acrylate wt%: 7.5, 15, 22.5%
- UV dose: 37, 105, 172 mJ/cm^2^

### Particle recovery and washing

Following polymerization, excess oil was removed by pipetting and PBS was added to the remaining emulsion. To destabilize the emulsion and transfer particles into the aqueous phase, a solution of 20% (v/v) perfluorooctanol (PFO; Sigma) in Novec 7500 was added and mixed until phase transfer occurred. Remaining oil was removed, and particles were washed two times with Novec 7500 to remove residual surfactant. Novec 7500 was then removed by pipetting, and residual oil was removed by washing three times with hexane (Sigma). Particles were subsequently washed three times with PBSP and stored in PBSP (PBS pH 7.4 with 0.1% Pluronic F-127).

### Protein porosity assay

All 36 particle formulations (biotinylated during fabrication) were incubated directly with 16.5 nM of one of the following streptavidin–fluorophore conjugates: streptavidin Alexa Fluor 555 (Invitrogen), streptavidin allophycocyanin (Molecular Probes), or streptavidin R-phycoerythrin (Jackson ImmunoResearch). For each formulation, 25 µL of particles were incubated with 100 µL fluorophore solution for one week at room temperature on a rotator, protected from light (foil-wrapped). A one-week incubation was used for all conditions. Although selected to approach high probe loading, fluorescence continued to increase beyond one week in the formulation evaluated in the time-course experiment (Supplementary Fig. 4); the measurements therefore represent accumulation after a standardized incubation period, not the saturated condition. Following incubation, particles were washed three times with 500 µL PBSP and analyzed. Fluorescence imaging settings were kept constant across conditions.

### Gold porosity assay

Particles fabricated at one UV dose (105 mJ/cm^2^; 12 formulations spanning 4 molecular weights × 3 wt%) were incubated for one week with streptavidin–gold nanoparticles (Cytodiagnostics): 5 nm, 10 nm, and 20 nm AuNPs at 3 OD. Incubations were performed at room temperature on a rotator and protected from light. Particles were washed three times with 500 µL PBSP and analyzed by brightfield imaging. Imaging settings were fixed across conditions.

### Immunoassay

Particles were first incubated with streptavidin (50 µg/mL, 1 h) to provide binding sites for biotinylated capture reagents. Particles were then incubated with biotinylated goat anti-mouse antibody at 0, 1, or 100 µg/mL (2 h), followed by incubation with a mouse anti-human antibody conjugated to 5 nm gold nanoparticles (3 OD, 1 week) on a rotator and protected from light. The 0 µg/mL biotinylated goat anti-mouse condition served as the negative control (no capture antibody present on the particles).

Color intensity was quantified by converting images to HSV color space and computing the mean HSV saturation within a particle mask, where the mask was generated using the same approach described in Supplementary Fig. 1.

### Data collection

Fluorescence microscopy images were collected using an inverted fluorescence microscope (Nikon Eclipse Ti2) with a 10× objective through a TRITC filter set. Exposure time (500 ms) and camera gain were held constant across all conditions. Brightfield/color images were collected using an inverted microscope equipped with a color camera (Nikon). Imaging settings were kept constant across conditions within each assay.

### Analysis

Microscopy images were analyzed in ImageJ using a masking strategy (Supplementary Fig. 1). Background intensity was computed as the mean of three particle-free background regions per imaging session. Particle-associated fluorescence was quantified as background-corrected intensity normalized by particle area. 70 particles were measured for each condition and error bars demonstrate standard error. For gold colorimetric analysis, images were converted to HSV color space and saturation was averaged within the particle mask as described above. 20 particles were measured for each condition.

### Swelling measurements

Dry particle diameter was measured on the Nikon microscope at the time of fabrication. Swollen particle diameter was derived from particle pixel area using the same particle masking approach described in Supplementary Fig. 1. Swelling ratio was calculated from the ratio of swollen to dry volumes assuming spherical geometry.

## Data availability

The data supporting this article have been included as part of the electronic submission. Raw data are available upon request

## Author contributions

Alyssa Arnheim (A.A.) outlined and wrote the manuscript, fabricated the particles, collected and analyzed the data herein, and created the figures. Ian Morales (I.M.) fabricated the particles, collected and analyzed the data herein, and edited the manuscript. Andrew Tran (A.T) collected and analyzed data and edited the manuscript. Dino Di Carlo (D.D.) contributed writing to the introduction and discussion sections, and edited the final manuscript and figures.

## Acknowledgements

The authors acknowledge the National Science Foundation PATHS-UP Engineering Research Center (Grant No. 1648451) for providing funding for this work.

