## Supplemental files for "Hydrogel formulation controls size-dependent accumulation of proteins and nanoparticles in PEG microparticles"

### Supplementary materials

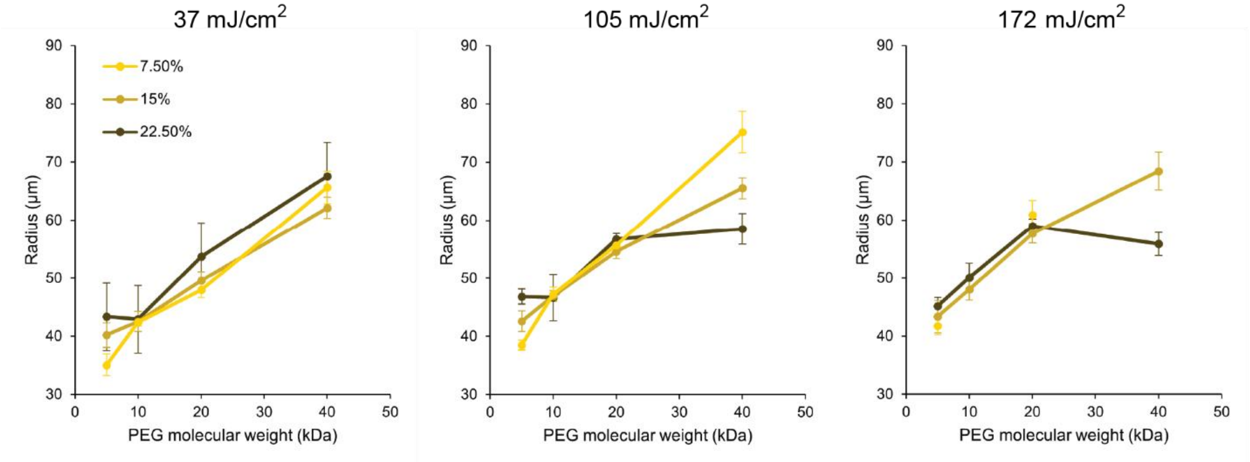


Supplementary Figure 1. Swollen radii of all fabricated particles as a function of PEG molecular weight. Radii were measured by converting pixel area of all particles measured back to radii.

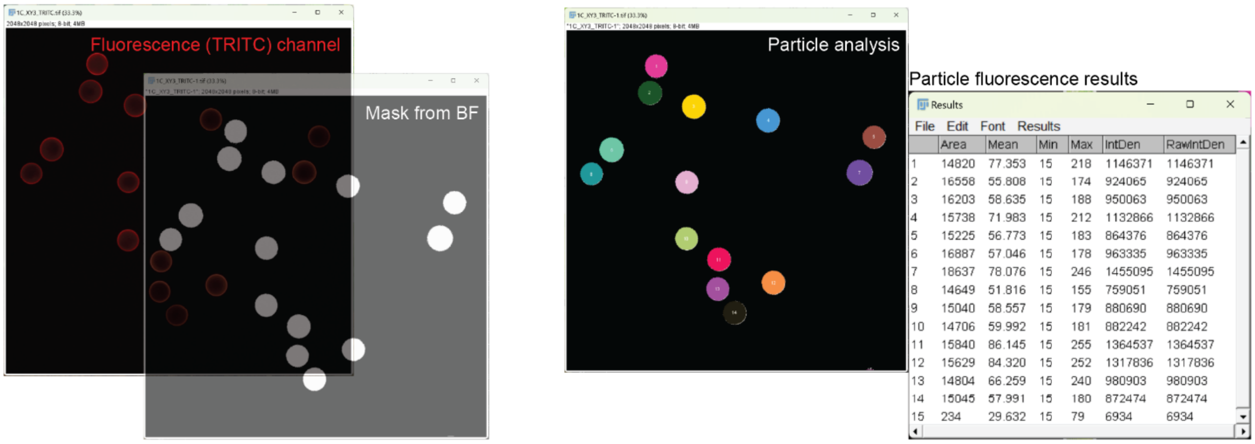


| | Area | Mean | Min | Max | IntDen | RawIntDen |
|---|---|---|---|---|---|---|
| 1 | 14820 | 77.353 | 15 | 218 | 1146371 | 1146371 |
| 2 | 16558 | 55.808 | 15 | 174 | 924065 | 924065 |
| 3 | 16203 | 58.635 | 15 | 188 | 950063 | 950063 |
| 4 | 15738 | 71.983 | 15 | 212 | 1132866 | 1132866 |
| 5 | 15225 | 56.773 | 15 | 183 | 864376 | 864376 |
| 6 | 16887 | 57.046 | 15 | 178 | 963335 | 963335 |
| 7 | 18637 | 78.076 | 15 | 246 | 1455095 | 1455095 |
| 8 | 14649 | 51.816 | 15 | 155 | 759051 | 759051 |
| 9 | 15040 | 58.557 | 15 | 179 | 880690 | 880690 |
| 10 | 14706 | 59.992 | 15 | 181 | 882242 | 882242 |
| 11 | 15840 | 86.145 | 15 | 255 | 1364537 | 1364537 |
| 12 | 15629 | 84.320 | 15 | 252 | 1317836 | 1317836 |
| 13 | 14804 | 66.259 | 15 | 240 | 980903 | 980903 |
| 14 | 15045 | 57.991 | 15 | 180 | 872474 | 872474 |
| 15 | 234 | 29.632 | 15 | 79 | 6934 | 6934 |

Supplementary Figure 2. Fluorescence signal intensity analysis pipeline. A particle mask generated from the brightfield image is projected onto the raw pixel intensities from the fluorescence channel of interest (FITC). Individual particle measurements are extracted.

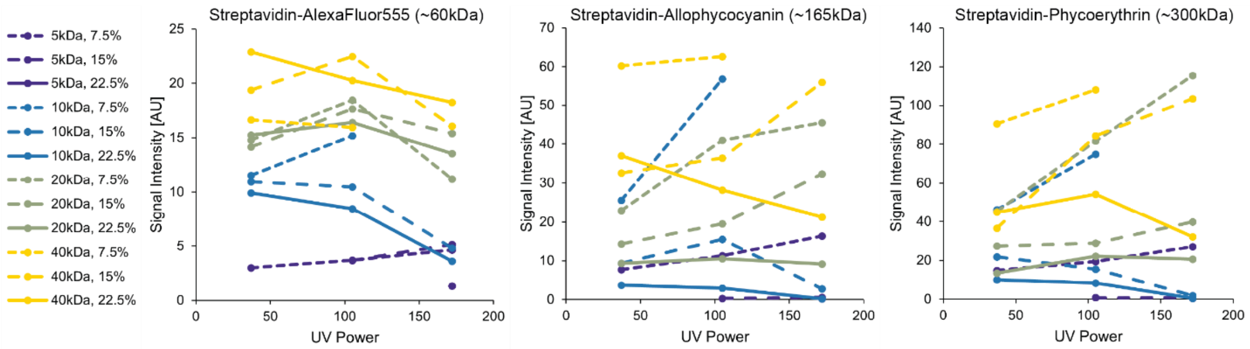

Supplementary Figure 3. Impact of UV dose on final particle-associated fluorescence intensity. Particle-associated fluorescence was measured after incubation with streptavidin–Alexa Fluor 555 (~60 kDa), streptavidin–allophycocyanin (~165 kDa), or streptavidin–phycoerythrin (~300 kDa) across PEG-acrylate formulations spanning nominal PEG-acrylate molecular weight (5–40 kDa) and PEG weight percent (7.5–22.5 wt%). Curves connect measurements obtained at three UV crosslinking doses (37, 105, and 172 mJ/$cm^2$). Line color indicates nominal PEG-acrylate molecular weight and line style indicates PEG wt%. Fluorescence intensity is reported as background-corrected signal normalized by particle area (see Methods), enabling comparisons across formulations with different particle sizes. Across probes, UV dose did not produce a consistent monotonic trend: depending on formulation, increasing UV dose sometimes increased and sometimes decreased the final signal, and UV-dependent changes were generally subtle relative to the strong effects of nominal PEG-acrylate molecular weight and PEG wt%.

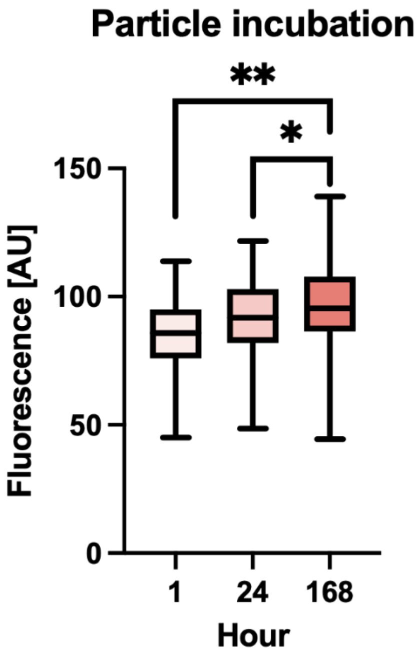


Supplementary Figure 4. Particle fluorescence over different incubation times. To determine how long particles should be incubated with fluorophore, we incubated 10 kDa 15% 105mJ/$cm^2$ particles with streptavidin-PE (10 µg/mL) for the incubation times listed, washed and measured. ** p <0.0001, * p 0.0002
